# Global analysis of protein turnover dynamics in single cells

**DOI:** 10.1101/2024.05.30.596745

**Authors:** Pierre Sabatier, Zilu Ye, Maico Lechner, Ulises H. Guzmán, Christian M. Beusch, Fabiana Izaguirre, Anjali Seth, Olga Gritsenko, Sergey Rodin, Karl-Henrik Grinnemo, Jesper V. Olsen

## Abstract

Even with recent improvements in sample preparation and instrumentation, single-cell proteomics (SCP) analyses mostly measure protein abundances, making the field unidimensional. In this study, we employ a pulsed stable isotope labeling by amino acids in cell culture (SILAC) approach to simultaneously evaluate protein abundance and turnover in single cells (SC-pSILAC). Using state-of-the-art SCP workflow, we demonstrated that two SILAC labels are detectable from ∼4000 proteins in single HeLa cells recapitulating known biology. We investigated drug effects on global and specific protein turnover in single cells and performed a large-scale time-series SC-pSILAC analysis of undirected differentiation of human induced pluripotent stem cells (iPSC) encompassing six sampling times over two months and analyzed >1000 cells. Abundance measurements highlighted cell-specific markers of stem cells and various organ-specific cell types. Protein turnover dynamics highlighted differentiation-specific co-regulation of core members of protein complexes with core histone turnover discriminating dividing and non-dividing cells with potential in stem cell and cancer research. Our study represents the most comprehensive SCP analysis to date, offering new insights into cellular diversity and pioneering functional measurements beyond protein abundance. This method distinguishes SCP from other single-cell omics approaches and enhances its scientific relevance in biological research in a multidimensional manner.

## Introduction

The stability of proteins defined by their turnover is crucial for their function and plays a regulating role in multiple cellular and biological mechanisms^1,2^. Protein turnover dynamics is particularly important in modulating signaling cascades and receptor recycling/degradation^3–5^ and thus is also involved in multiple diseases including cancers^6^. A famous example of this is the activity of p53, often called the guardian of the genome, which is regulated by degradation^7–9^. In addition, protein turnover is also exploited by pharma/biotech in proteolysis targeting chimera (PROTAC) drugs^10,11^ and can provide information on post-translational modifications (PTM) dynamic^12,13^ and thus represent a key aspect of protein chemistry. One of the unique features of mass spectrometry (MS)-based proteomics is that it is currently the only technology allowing unbiased and system-wide measurements of protein synthesis, degradation and turnover rates, thanks to pulsed Stable Isotope Labeling by Amino acids in Cell culture (pSILAC)^14–18^. However, pSILAC is only applicable to bulk analysis of cell lines or tissues, and can only estimate average protein turnover rate of a cell population, which often includes different cell-types and cells undergoing all the various stages of the cell cycle. Advances in single-cell (SC) analysis and in particular from SC transcriptomics studies have shown that cell heterogeneity in cell lines and tissues is higher than what was assumed prior to the advent of these technologies. This high heterogeneity in cell populations is also highlighted by mass spectrometry-based single-cell proteomics (SCP) for which powerful methods that have emerged recently^19–24^. Microscopy-based analyses showed that protein turnover rate can be different from one cell to another both globally and for individual proteins and studying protein turnover in SC provides a new layer of functional insight in cellular and molecular biology^25^. However, such studies are either limited to untargeted and/or to a low number of specific proteins only, and while pSILAC analysis in SCP would provide a system-wide view of protein turnover, it is currently inexistent because of technical challenges.

SILAC-based MS analysis has traditionally been performed in data-dependent acquisition (DDA) mode with full-scan MS^1^-based quantification. Conversely, label-free SCP analyses greatly benefit from the higher sensitivity offered by DIA methods with MS^2^-based quantification. SILAC relies on metabolic incorporation of isotopically labeled amino acids into proteins. For trypsin digestion-based proteomics, lysine and arginine are the preferred SILAC amino acids with a mass difference of 8 and 10 Da at most, for the heavy versions compared to the light SILAC versions. However, for multiply-charged peptide ions such a mass spacing is smaller than most conventional wide-window DIA isolation window schemes. Therefore, individual DIA-MS^2^ spectra most often contain fragment ions from both heavy and light SILAC labeled versions of a peptide. In MS^2^ spectra of tryptic peptides derived from arginine and lysine-labeled SILAC peptides, only y-ions are different in mass, whereas N-terminal fragments have identical masses including a- and b-ion series^26,27^. This renders MS^2^–based quantification of SILAC data challenging in DIA mode, and consequently MS^1^ quantification is preferred since it is more straightforward, reproducible and accurate^27^. Advances in computational proteomics software^26,27^ including popular DIA search engines like Spectronaut and DIA-NN^28^ allow mitigating this effect and to exclude the shared fragment ions. While this works well for bulk proteomics, it is an issue for SCP analysis as the average intensities of in MS^2^ spectra are very low resulting in decreased quantification accuracy, particularly in a SC-pSILAC experiment in which the general peptide abundance is lower than standard SCP as the MS signal is split between the two SILAC-labeled precursors. Here, we capitalized on the recent developments in high-sensitivity MS instrumentation and DIA approaches^27^, particularly for multiplexed DIA or plex-DIA^29–31^ with the narrow-window (n)DIA enabled by the Orbitrap Astral^32^ as well as improvements in SCP sample preparation^31,33^ increasing the sensitivity of the analysis, to perform single-cell pSILAC. We first applied pSILAC to minute cell numbers and SCs and demonstrated that we can detect and quantify both light and heavy labels and that the corresponding SILAC ratios in SCs are in accordance with those from multiple cells and bulk analysis, providing functional information on protein turnover at the SC level. We furthermore showed that by treating cells with the proteasome-inhibitor bortezomib or the protein translation-inhibitor cycloheximide, turnover analysis reflected the expected disturbances in protein synthesis and degradation in single cells. This strategy effectively highlights the specific differences observed in protein turnover dynamics by treatment with different drugs and concentrations compared to protein abundance measurements. Finally, we applied SC-pSILAC to a large-scale analysis of undirected differentiation induced pluripotent stem cells (iPSCs) through embryoid body (EB) induction collecting data over two months of differentiation and analyzing the proteome turnover dynamics in more than one thousand individual cells. Exploring this rich dataset highlighted cell and stage-specific variations in protein expression and turnover.

## Results

### Pulsed (p)SILAC is amenable to SCP analysis

To test whether we could obtain information on protein turnover in single cells, we used a pSILAC approach by switching cells grown in a growth medium containing non-isotopically labeled amino acids (light SILAC) to a medium containing heavy SILAC L-lysine-^13^C_6_^15^N_2_ and L-arginine-^13^C_6_^15^N_4_ resulting in tryptic peptides with mass differences of +8 and +10 Da, respectively. We analyzed bulk, 10-cell pools and single HeLa cells (Figure 1a). After 18 hours post SILAC label switch, corresponding to about one cell cycle doubling and therefore on average half incorporation of the new heavy SILAC label in term of total intensity, cells were detached, trypsinized to make single cell suspensions and sorted using the cellenONE into a label-free (LF-48) proteoCHIP. Proteins were extracted and digested using a master mix consisting of 0.2% n-Dodecyl-β-D-Maltoside (DDM), 100 mM TEAB, 10 ng/µl lysyl endopeptidase and 20 ng/µl trypsin. Resulting peptide mixtures were loaded onto Evotips and analyzed using a whisper flow gradient at 40SPD on an Evosep One LC system connected to an Orbitrap Astral mass spectrometer operated in narrow-window DIA mode with Orbitrap full scans and parallel acquisition of 100Hz DIA-MS^2^ by the Astral analyzer (Figure 1a). Multiple pairs of light and heavy SILAC labels were visible in full-scan MS^1^ spectra, corresponding to incorporation of heavy lysine and arginine isotopic variants separated by either 8 or 10 Da (Figure 1b). This demonstrated that the MS^1^ signal was sufficiently high for the Orbitrap analyzer to detect both labels. We processed the raw MS files using directDIA searches in Spectronaut adding the SILAC modifications as a second channel for quantification and excluding shared fragments in MS^2^. On average, 4288 and 3041 proteins (Figure 1c) and 39,169 and 22,494 peptides (Figure 1d) were quantified in 10 and single Hela cells, respectively. Both labels were quantified in 3737 and 2781 proteins (Figure 1e) in 10 and single HeLa cells, demonstrating that pSILAC is possible in SC.

**Figure 1.**
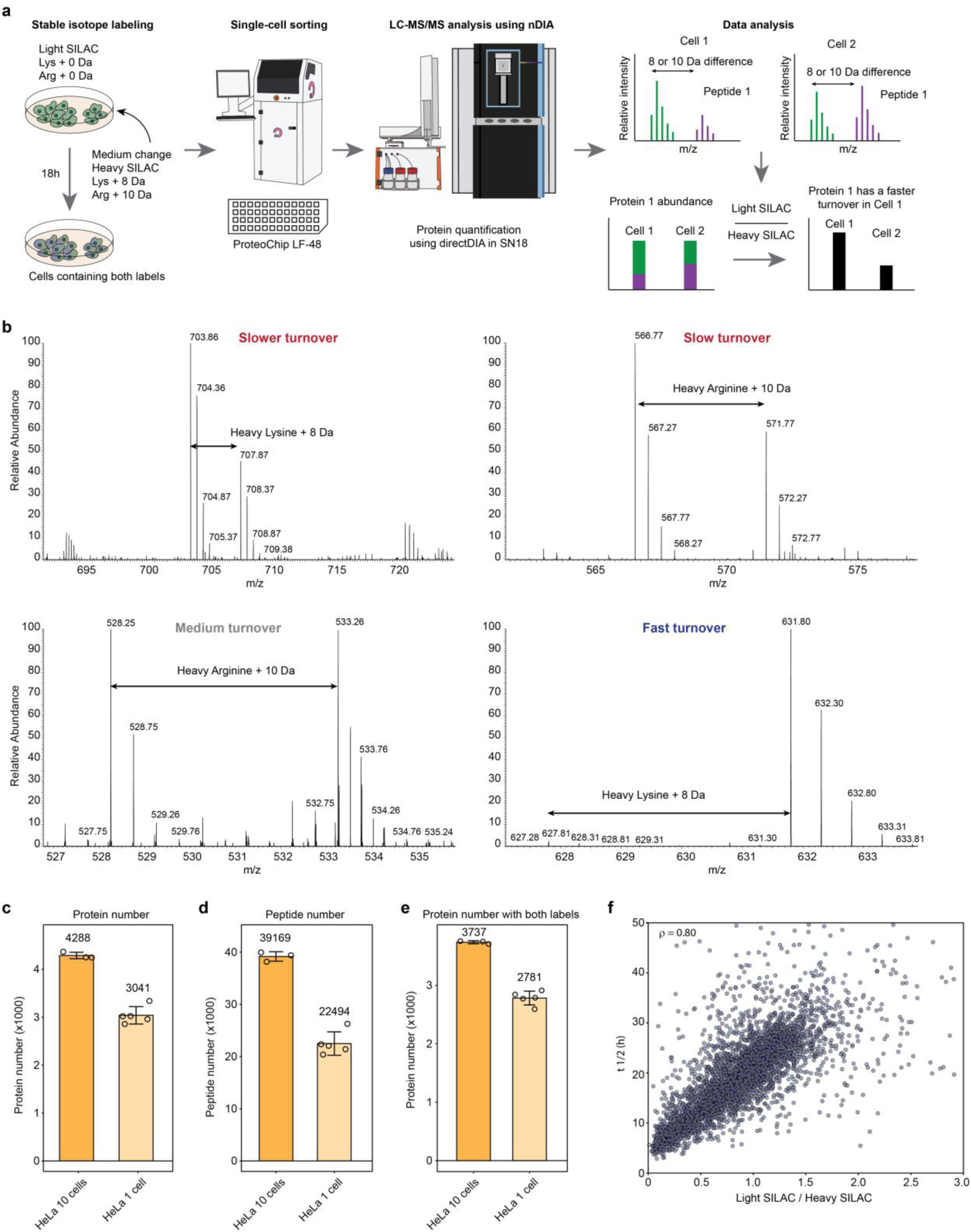
Single-cell pulsed SILAC (SC-pSILAC). (**a**) Workflow of the pSILAC experiment. (**b**) Representative Orbitrap full-scan MS spectra highlighting charge +2 peptides harboring light (+0 Da) and heavy (+8 Da for lysine or +10 Da for arginine) SILAC labels. (**c**), (**d**), (**e**) Number of proteins, peptides, and proteins harboring the two SILAC labels, respectively, identified in 10 HeLa cells and single HeLa cells. The mean is indicated. Error bars represent ± the standard deviation of the mean. (**f**) The ratio of light versus heavy SILAC in human foreskin fibroblasts (hFF) after a single 24 h pulse and estimated protein half-life in hFF from a time series measurement. Spearman correlation between the two measurements is displayed on the plot. n=3 for 10 HeLa cells and n=5 for single HeLa. n=3 for hFF.

### SC-pSILAC estimates protein turnover in single cells and provides complementary information to abundance measurement

Since our goal was to obtain information about protein turnover, we utilized the ratio of the light SILAC to heavy SILAC labels (relative turnover^34^), where light SILAC-labeled proteins denote the pre-existing pool present at the time of the switch in growth media, while heavy SILAC-labeled proteins represent newly synthesized proteins post-switch. Therefore, proteins exhibiting a high light-to-heavy (L/H) ratio indicate a slow turnover, whereas proteins with low L/H ratio suggest a fast turnover. However, the limitation of such measurement in single cells is that we can only obtain a single time point. Thus, to demonstrate that a single sampling time can still yield information on protein turnover, we performed a time-series bulk analysis of SILAC-label incorporation in a homogeneous human foreskin (hFF) cell line. We calculated protein half-life (T_½_)^35,36^ for each protein and plotted it against the L/H SILAC ratio from an independent single-point SILAC measurement after 24 h of labeling. The Spearman correlation between these two independent biological measurements was 0.8, indicating that our single-point SILAC approach provides insights into protein turnover and half-life (Figure 1f).

To confirm that biological information about specific protein turnover is maintained even at the single cell level, we performed unsupervised hierarchical clustering of the SILAC ratio of bulk, 10 and single HeLa cells data and compared the protein clusters showing fast or slow turnover and performed gene ontology (GO) enrichment analysis (Supplementary Figure 1a). Proteins showing faster or slower turnover in single cells mostly group in the same clusters as bulk and 10 cells with more extreme ratios in single cells likely due to cell heterogeneity or lower measurement accuracy. Clusters containing proteins with fast turnover encompassed proteins linked to cell division while clusters containing proteins with slower turnover were enriched for ribosomal proteins and histones among other pathways (Supplementary Figure 1a). Such protein turnover characteristics have been reported previously^15^ and proteins involved in cell division and in the cell cycle are known to be short-lived while proteins in larger complexes are usually more stable and have a slower turnover, which is also highlighted in our dataset (Supplementary Figure 1b). This result demonstrate that biological information is preserved and can be obtained from pSILAC analysis of single cells as well as in few cells as the measurements are globally in accordance with bulk analysis.

Since the information obtained through the light and heavy SILAC pairs is complementary to protein abundance measurements, we investigated whether they could also differentiate cells on principal component analysis (PCA). Individual pSILAC-labeled HeLa and HEK 293T cells separated clearly on PCA using the normalized sum of both labels as a proxy for abundance measurement (Supplementary Figure 1c). Similarly, they differentiated well in PCA using the ratio of the two labels, indicating that both measurements can effectively distinguish between different cell populations (Supplementary Figure 1d).

### Proteasome and translation inhibitors disrupt protein turnover in single cells

To demonstrate to ability to monitor protein turnover at the single cells level, we analyzed HeLa cells treated with 1 and 10 µM of the proteasome inhibitor bortezomib and with 2 and 20 µM of the protein translation inhibitor cycloheximide for 18 hours and compared them to non-treated cells (Figure 2a). To verify that the treatments had a global effect on the proteome of the cells, we plotted the total protein intensities for each sample and condition corresponding to 10 and single HeLa cells (Figure 2b). The overall protein intensities indicated minimal change in the light label globally, with slightly higher intensities observed for bortezomib-treated cells compared to the control group. Conversely, the heavy label exhibited a concentration-dependent decrease, particularly evident with cycloheximide treatment, indicating inhibition of new protein synthesis as expected. Consequently, all treatments affected global protein turnover.

**Figure 2.**
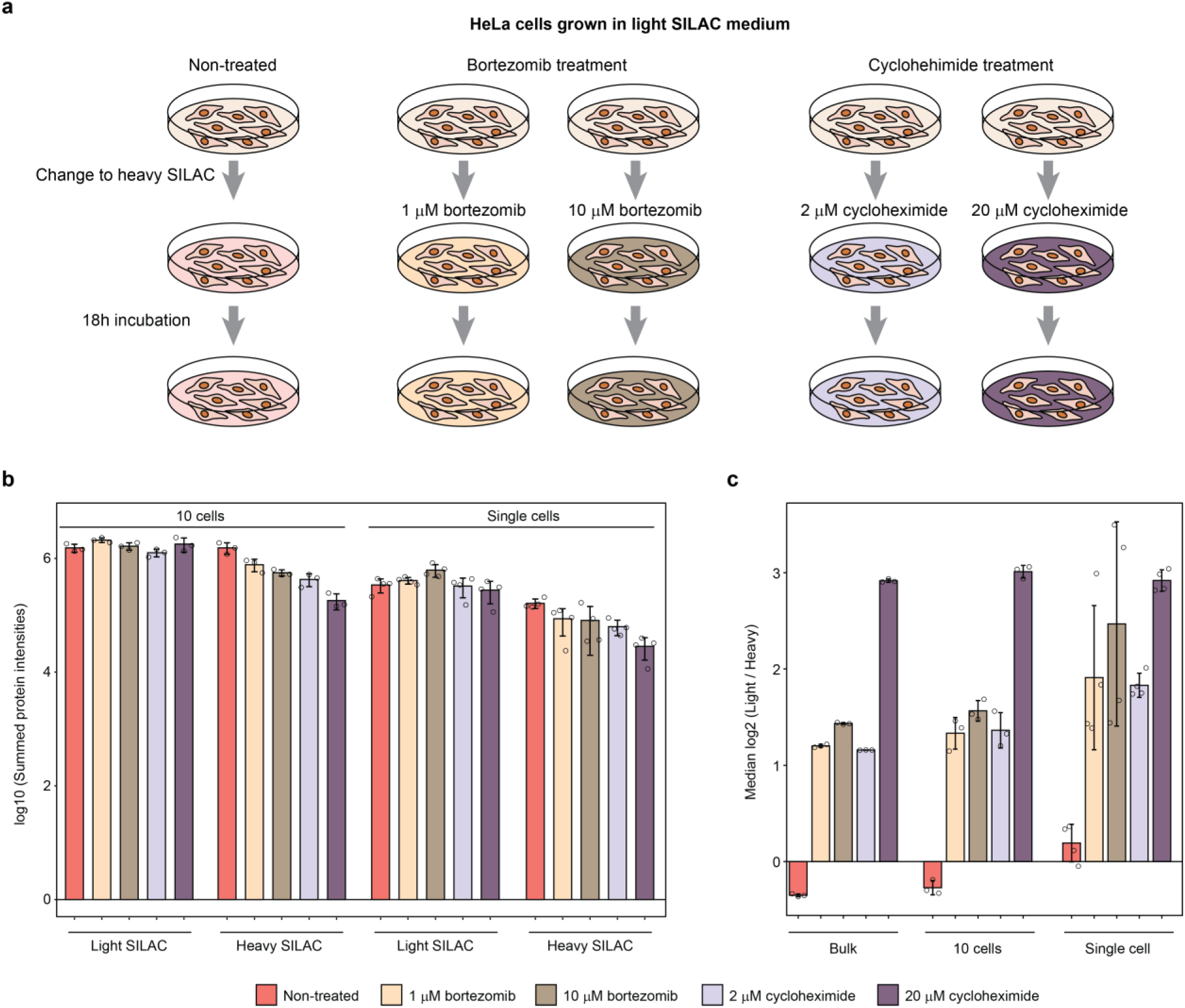
SC-pSILAC of HeLa cells treated with bortezomib and cycloheximide. (**a**) Workflow of the treatment and pSILAC. (**b**) log10 of the summed protein intensities in 10 Hela cells and single HeLa cells for the light and heavy SILAC labels for each treatment, namely non-treated, treated with 1 µM, 10 µM bortezomib and 2 µM and 20 µM cycloheximide. (**c**) Median of the relative turnover of light divided by heavy for bulk, 10 and single HeLa cells. Error bars represent ± the standard deviation of the mean. n=3 bulk, n=3 10 cells, n=4 single cells for each treatment.

To verify that similar alterations are observed at the single-cell level as in bulk, we next plotted the median protein relative turnover (L/H) of each treatment across bulk, 10 and single HeLa cells. All treatments significantly increased the turnover compared to the non-treated samples and revealed similar trends with both bortezomib and cycloheximide showing concentration-dependent increase in the L/H SILAC ratio with cycloheximide exhibiting the highest median ratio. The fact that the calculated median relative turnover was similar level between bulk, 10 and single HeLa cells further reinforced the reliability of our methodology in capturing protein turnover information at the single-cell level. This offers a promising avenue for further research into the cellular responses to different biochemical treatments and their implications on protein dynamics. Additionally, we found that SC-pSILAC helped distinguishing treatments altering protein turnover better than abundance measurements and is less sensitive to sample size (Supplementary Figure 2).

### SC-pSILAC distinguishes protein degradation inhibitor from synthesis inhibitor

Next, we compared the relative abundance level of light and heavy SILAC labels between non-treated samples, bortezomib and cycloheximide treatments. While the SILAC ratio can provide an estimate for protein turnover, it cannot distinguish increase or decrease in protein synthesis and can only conclude on the amplitude of the effect on protein turnover. Therefore, we analyzed the light and heavy SILAC labels separately with the light SILAC providing information on degradation only, while the heavy SILAC contains a mixed readout of synthesis and degradation. We normalized the labels by the corresponding labels in the non-treated cell group. Since the level of degradation in the light SILAC label is expected to be the same as in the heavy SILAC label, the additional changes in heavy label can be interpreted as changes in protein synthesis. Consequently, this approach enables us to capture a nearly comprehensive view of the turnover dynamics of a particular protein. When analyzing the mean ratio for light SILAC label between treated and control, bortezomib-treated sample group showed significant increase compared to control, as does cycloheximide treatment but to a lower extent (Figure 3a). Conversely, the situation was reversed for the heavy SILAC label for which the ratio decreases massively for cycloheximide particularly at 20 µM highlighting inhibition of protein synthesis. Interestingly, the ratios are much lower in bortezomib for the heavy SILAC label than they are for the light SILAC label showing that translation inhibition also occurs upon bortezomib treatment as a response to proteasome inhibition. Additionally, bortezomib clearly showed a concentration-dependent effect on protein degradation as the distribution of the ratios of the light SILAC label is significantly higher at 10 µM than at 1 µM, while cycloheximide did not show significant changes between 20 and 2 µM on the light SILAC label. Conversely, for the heavy SILAC label, cycloheximide showed a large difference between 2 and 20 µM while it was barely significant (p=0.03) between the two concentrations of bortezomib. Even though both drug treatments have an effect on both protein synthesis and degradation, their mode of action is quite different as highlighted by the analysis. This analysis also showed that we can potentially distinguish effects that arise from inhibition of degradation or synthesis at the SC level.

**Figure 3.**
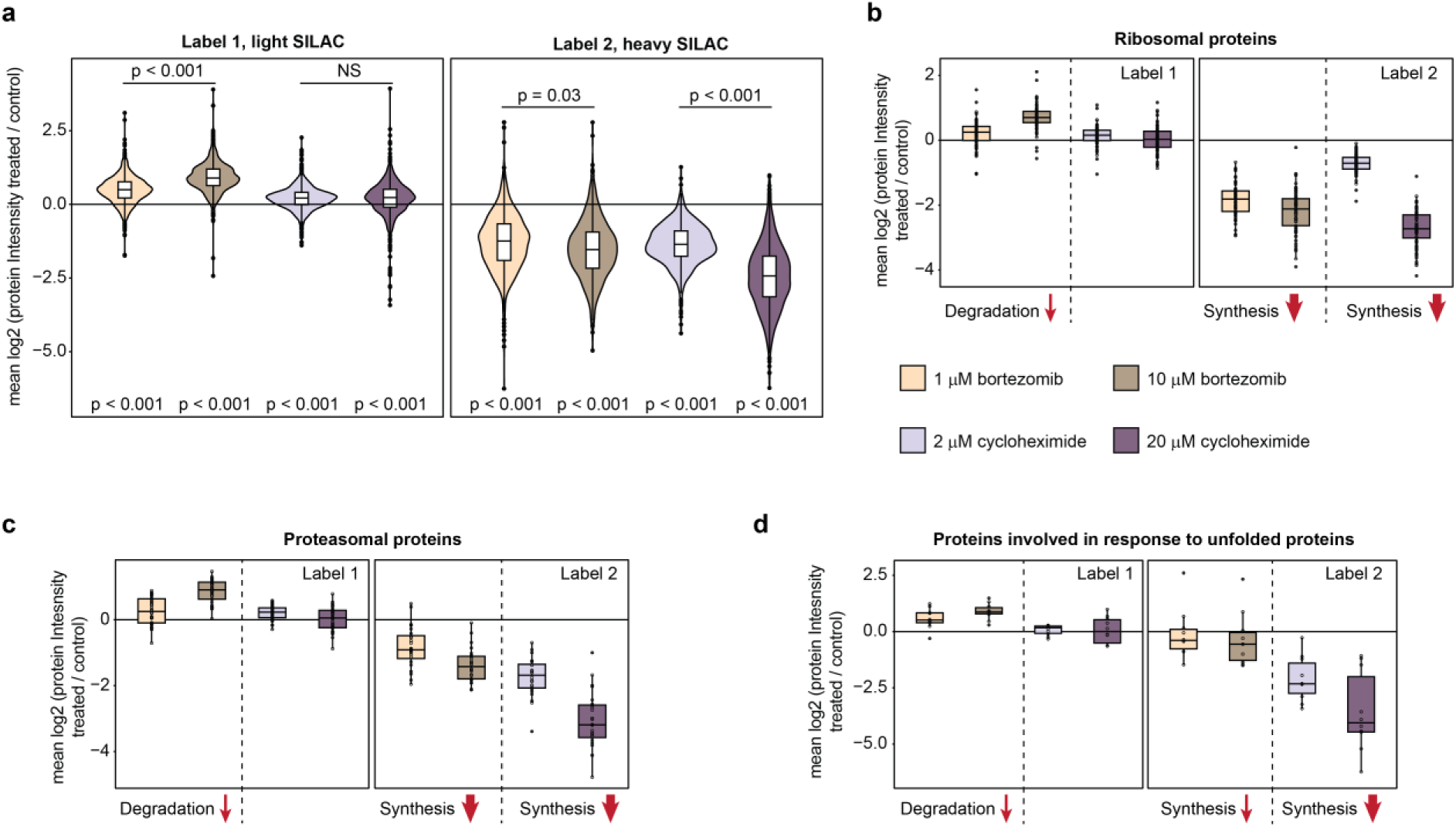
SC-pSILAC highlights differences in bortezomib and cycloheximide treatment. (**a**) Mean log2-transformed MS signal intensities of light and heavy SILAC labels in 1 µM, 10 µM bortezomib and 2 µM and 20 µM cycloheximide treatments relative to control (non-treated). (**b**), (**c**), (**d**) Same measurements highlighting protein groups that were enriched in difference between, 10 µM bortezomib and 20 µM cycloheximide and non-treated control corresponding to ribosomal proteins, proteasomal proteins and proteins involved in response to unfolded proteins. Red arrows represent increase or decrease in protein synthesis or degradation. The GO enrichment was performed using DAVID. P-values were calculated using a two-sided Student t-test. Horizontal line in the boxplots represent the median, 25th and 75th percentiles and whiskers represent measurements to the 5th and 95th percentiles. n=4 for each treatment.

Lastly, we plotted the distribution of protein ratios of each treatment against non-treated cells for the groups of proteins that had significantly altered turnover and showed GO enrichment in ribosomal and proteasomal proteins and proteins involved in response to unfolded proteins (Figures 3b-d). The results were generally in accordance with what was observed with all quantified proteins. However, for bortezomib treatment we observed a much higher decrease in ribosomal protein level on heavy SILAC label and only a slight decrease in proteins involved in response to unfolded proteins. This is likely a feedback mechanism as cells with inhibited proteasome activity by bortezomib tend to shutdown protein synthesis and increase the response to unfolded proteins arising from the lack of degradation^37,38^. Taken together, our results demonstrate that we can detect differences in turnover, degradation and synthesis both at the proteome and individual protein level in SCs.

Finally, we sought to investigate whether we could detect treatment-related changes in abundance and turnover aimed at a specific protein. Therefore, we treated HeLa cells with 10 µM of the glycogen synthase kinase 3 (GSK3) inhibitor CHIR99021 and performed SC-pSILAC analysis. Inhibition of GSK3 prevents phosphorylation and subsequent degradation of β-catenin (CTNB1), which instead accumulates in the cytoplasm leading to its translocation to the nucleus and activation of the canonical Wnt pathway transcriptional program^39^ (Figure 4a). We were thus expecting a change in CTNB1 abundance and/or turnover. Since analytical depth is crucial to study specific effector proteins at the SC level, we employed the newly developed Evo96 proteoCHIP allowing direct transfer to Evotips^33^ and quantified more than 4000 proteins on average with both SILAC labels (Supplementary figure 3). We observed a significant increase in CTNB1 abundance and a faster turnover compared to the DMSO-treated control (Figures 4b and c). Interestingly, variation in the magnitude of abundance and turnover changes between individual cells also highlighted single cell heterogeneity in response to the treatment. When plotting the ratios of protein abundances and relative turnover in CHIR99021 treatment against control, CTNB1 stood out as a strong outlier (Figure 4d). The Pearson correlation between the two measurement types was -0.27 indicating their lack of correlation and thus highlighting their complementary nature. Finally, CTNB1 ranked first and third according to the ratio of abundance and relative turnover of CHIR99021 treatment against control, respectively (Figures 4e and f). This study of GSK3 inhibition demonstrated that the combined measurements of abundance and protein turnover in SCs can pinpoint specific proteins affected by drug treatments and reflect heterogeneity in the response from individual cells.

**Figure 4.**
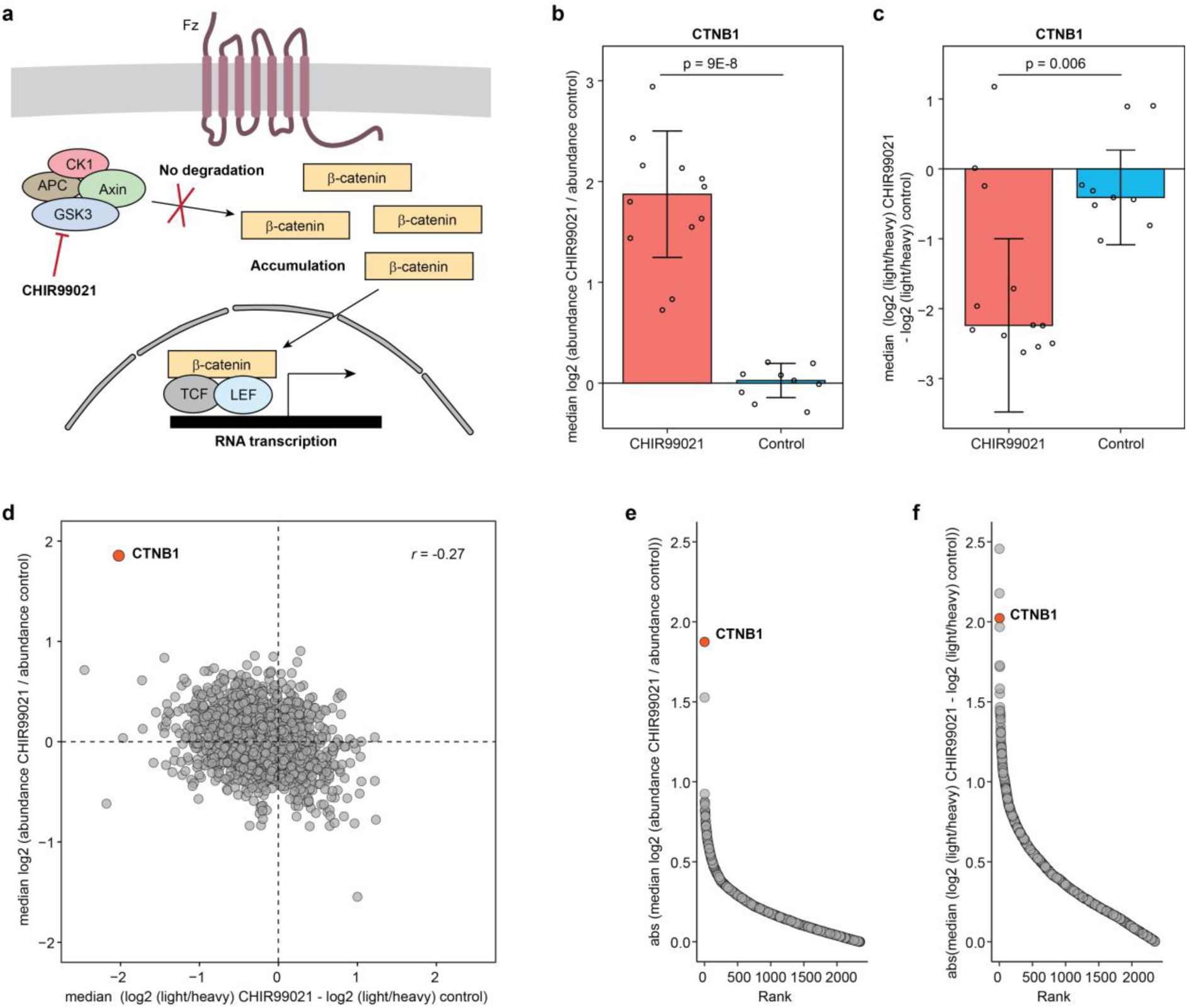
SC-pSILAC of HeLa cells treated with the GSK3 inhibitor CHIR99021 showing accumulation of β-catenin. (**a**) Inhibition of GSK3 prevents phosphorylation of β-catenin (CTNB1) and subsequent proteasome degradation, and instead promotes accumulation and migration to the nucleus to activate transcription. (**b**), (**c**) Median CTNB1 abundance and median relative turnover in CHIR99021 treatment compared to DMSO-treated control, respectively. P-values were calculated using a two-sided Student t-test. Error bars represent ± the standard deviation of the mean. n=12 for CHIR99021 treatment and n=9 for DMSO-treated control. (**d**) A scatter plot of the median relative turnover (x-axis) and median abundance (y-axis) in CHIR99021 treatment compared to DMSO-treated control, β-catenin is highlighted in orange. (**e**), (**f**) Protein ranking based on the absolute value of the median abundance and median relative turnover in CHIR99021 treatment compared to DMSO-treated control, respectively. CTNB1 is highlighted in orange.

### SC-pSILAC analysis of undirected stem cell differentiation

To study the dynamics of protein abundance and turnover within a biologically more complex system, we differentiated human induced pluripotent stem cells (iPSCs) through embryoid body (EB) induction. Subsequently, we performed SC-pSILAC analysis at six different time points over two months, alongside to two independent samplings of the iPSCs (Figure 5a). Plated EBs can differentiate into a myriad of cell types offering a great model to study the dynamics of protein abundance and turnover during the differentiation process, with cells varying in size, protein expression, cell state and division rate. We recorded protein abundance and turnover in more than 1100 single cells with at least 100 cells per conditions and identified more than 6500 proteins in the whole dataset with up to about 4000 proteins quantified with both labels in some large EB cells (Figure 5b). To first address potential biases in SILAC incorporation rates within the cell cultures, we incorporated a control step involving bulk measurements of hFF at each sampling time. In brief, we prepared a frozen stock of hFF from the same passage, thawed and plated a vial of those cells 3 days prior to each sampling point, light SILAC growth medium was switched to heavy SILAC at the same time as for the iPSCs and EB cells, and samples were collected concurrently with the single-cell samples and prepared for bulk proteomics analysis. Notably, the coefficient of variation of the median SILAC ratios in hFF over two months and samplings was 8.1%, indicating highly stable and consistent SILAC incorporation throughout our experiment (Supplementary Figure 4).

**Figure 5.**
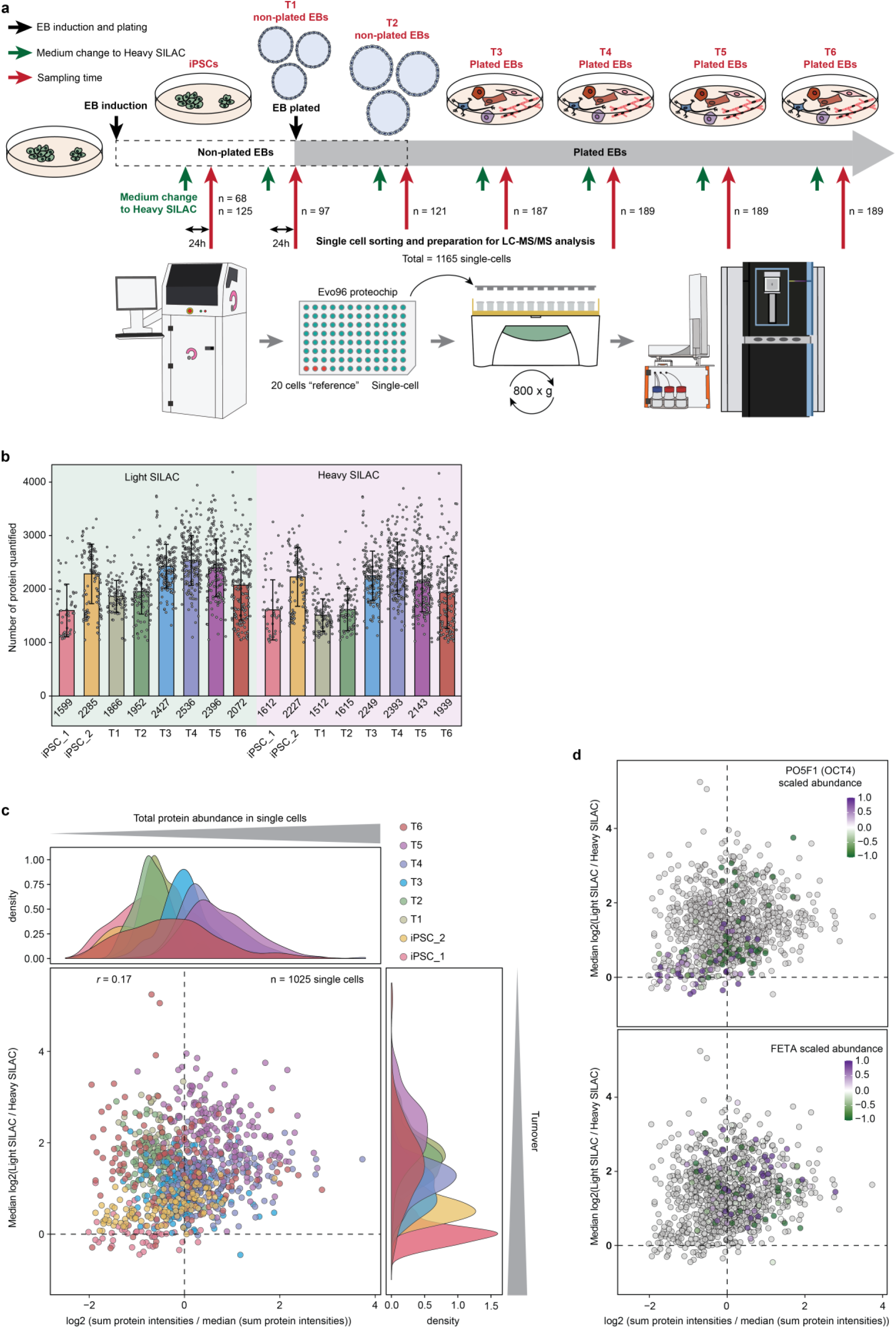
SC-pSILAC analysis of undirected differentiation of iPSCs through EB induction. (**a**) Workflow describing differentiation, pulsed SILAC and sampling timing as well as the number of cells and the sample preparation method used. (**b**) Number of protein quantified with light and heavy SILAC labels in every cell analyzed. For consistency, cells having less than 1000 proteins quantified with any of the labels were excluded from the analysis resulting in 1025 remaining cells. Cells were colored according to their sample group (**c**) Scatter plot of log 2 summed protein intensities (light + heavy SILAC) (x-axis) scaled by the median value among all cells versus the median log2 relative turnover (light / heavy) from every protein in all cells considered in the analysis (y-axis). Top and left represent the distribution of values of x and y-axis, respectively. Cells were colored according to their sample group. (**d**) Same scatter plot as in (**c**) but the coloring is according to OCT4 (top) and FETA (bottom) scaled abundance. Grey dots represent missing value.

The iPSCs and EB cells displayed a broad range of total protein intensities reflecting differences in cell size and of median relative turnover with the later matching expected fast cell dividing rates in iPSCs and slower dividing rates in differentiating EBs, with fast and slower incorporation of heavy SILAC, respectively (Figure 5c). Importantly, there was no apparent correlation between the two measurements (Figure 5c) confirming that they are orthogonal. Since we obtained abundance measurement alongside turnover by using the sum of the SILAC abundance, we first investigated the distribution of expression of specific cell-type markers. iPSCs expressed OCT4 (PO5F1) as expected and it was not present in most of the EBs cells while markers such as alpha fetoprotein (FETA) involved in fetal liver development were mainly co-expressed in few cells from EBs demonstrating that subsets of cell populations are present in our dataset (Figure 5d). The presence of OCT4 and FETA was confirmed by microscopy analysis along with other markers of the three embryonic germ layers in EB cells (ACTA, NEST, OTX2, SALL4 and TBB3) (Supplementary Figure 5). We also recorded the presence of beating cardiomyocytes (Supplementary media 1) and neuronal cells. Thus, the abundance of several other markers of various cell types was recorded even with lower analytical depth as compared to our established label-free analysis^33^ (Figure 6a) further demonstrating the wide range of cell types present in the dataset. This analysis enabled us to pinpoint some different cell type subsets such as iPSCs having high abundance of OCT4; hepatic cell lineages with FETA, FIBA, FIBG, HEMO EST1; melanocytes expressing PMEL; skeletal muscle cells with ACTS, MYL1; smooth muscle cells with ACTA; MYL4 for the cardiac cell lineage; and NEUM, TBB3, MTAP2, NFM, PEDF; for the neuronal lineage among others. Some clusters of cells harbored multiple markers allowing their identification as shown in Figure 6a, which further confirms the high heterogeneity of cell types in our dataset and that our approach can monitor the expression specific cell lineage markers differentiating between cell types.

**Figure 6.**
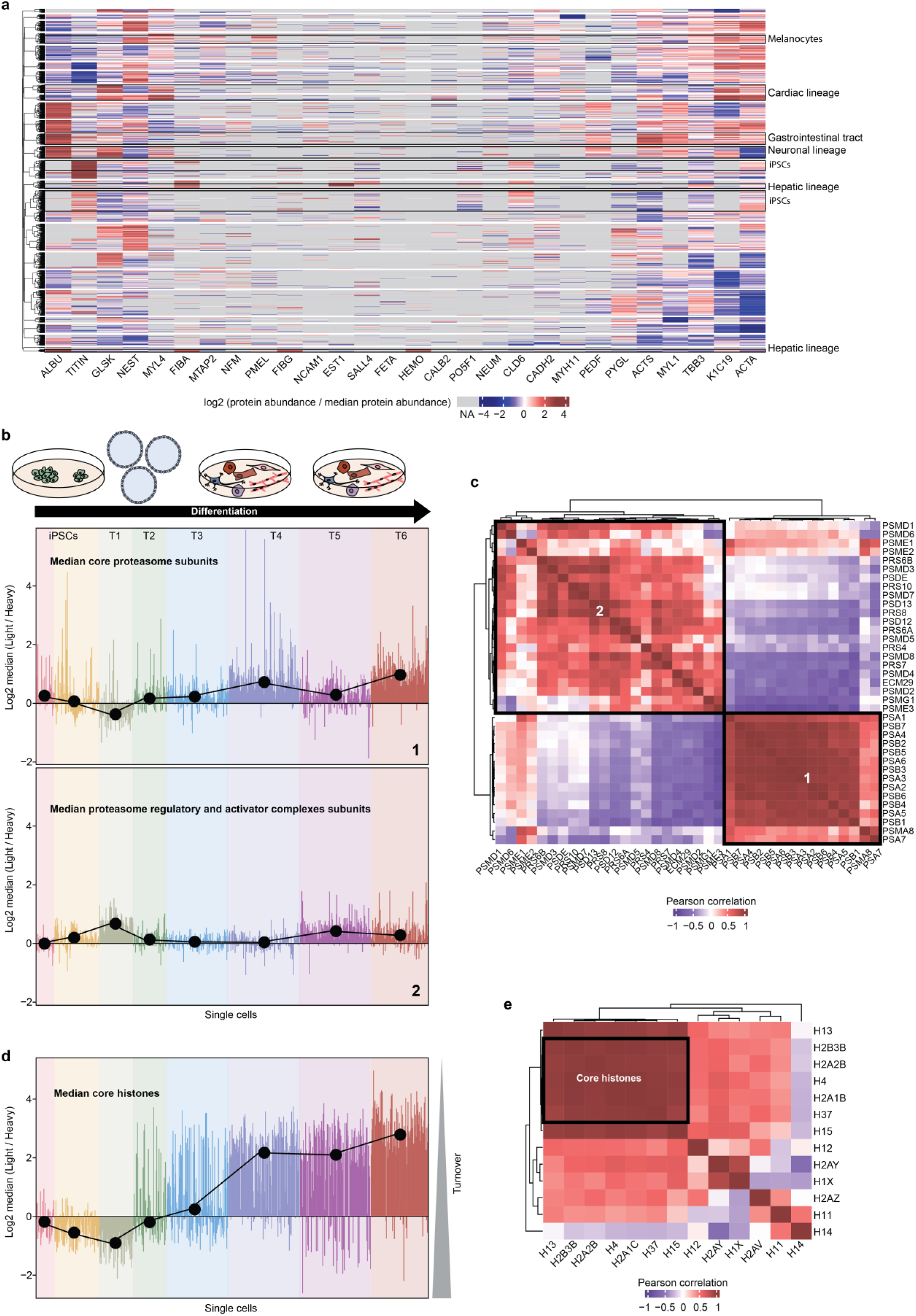
Cell-type specific markers expression and co-regulation of protein turnover in complexes. (**a**) Unsupervised hierarchical clustering using canberra and ward.D2 methods of proteins log2-transformed MS signal abundance scaled by the median abundance in all sample. (**b**) Median normalized relative turnover of core proteasome subunit (top) and of the regulatory and activators complexes (bottom) in every cell where these proteins were detected. The black dots represent the median value at each time point. (**c**) Unsupervised hierarchical clustering using canberra and ward.D2 methods of the Pearson correlation of the relative turnover of each proteasome subunit detected in the dataset, the median value at each time point was used to calculate the correlation. (**d**) Same as (**b)** with core histone proteins. (**e**) Same as (**c**) for histone proteins.

To compare the relative turnover of specific proteins between cells and time points, we normalized the L/H ratios. We then performed GO term enrichment analysis of the normalized median relative turnover of proteins for the 500 proteins with highest and lowest L/H SILAC ratios within each time point. Proteins with fast turnover were as expected enriched for pathways associated with cell cycle and cell division (Supplementary Figure 6). This was particularly evident for iPSCs along with pathways related to embryo development while fast turnover proteins were associated with markers of later stages of embryogenesis and cell differentiation, particularly in the formation of the three germ layers and of more mature tissues in later time points in differentiated cells. Proteins associated with higher L/H SILAC ratios with slow turnover were involved in pathways related to metabolism and ribosome assembly in iPSCs and progressively more involved in chromatin-related pathways as cells differentiate, highlighting the importance of chromatin regulation during differentiation (Supplementary Figure 7). Proteins with fast turnover are related to a wider variety of pathways comprising more specific features of differentiation than proteins with slow turnover suggesting that proteins with fast turnover can highlight cell-specific feature better during differentiation. Taken together, our data provides a timeline of turnover dynamics during differentiation.

### Turnover co-regulation of core members of protein complexes

Next, we investigated whether protein turnover dynamics analysis could be used to identify core members of protein complexes. To do this, we plotted the median relative turnover of the core subunits of the proteasome in each single cell analyzed as well as the proteasome activator and regulatory complexes (Figure 6b). The two proteasomal protein types showed different trends along the time of differentiation. The correlation of each subunit highlights a strong correlation between members of the core proteasome complex while there was little correlation with members of the activator and regulatory complex, which in turn correlated well with each other (Figure 6c). This demonstrated that turnover measurement can be used to study protein complex co-regulation in single cells and pinpoint core members of complexes.

We then performed a similar analysis for histones and plotted the median turnover for the core histones in each cell (Figure 6d). We observed a gradual decrease in protein turnover from iPSCs to time point 6 matching the decrease in cell dividing rate due to terminal differentiation and contact inhibition. We have previously demonstrated that differentiated cells have more stable histones due to increased compaction of their chromatin compared to pluripotent stem cells, which have a loser chromatin^40^. Our observation of histone turnover is in line with these observation. We then calculated the correlation of the median turnover of all histone proteins detected in more than half the cells present in the dataset (Figure 6e). Members of the core histone complex H2A3B, H2A2B, H37 and H4 clustered together with high correlation while they did not correlate with the non-conventional histones H2AY and H2AZ. Core histones also correlated highly with linker histones H13 and H15 showing that they are co-regulated while H11, H12 and H14 did not correlate. Interestingly, the abundance of histones did not correlate as well when considering the population median suggesting more variation in individual cells with the absence of a clear trend while histone turnover is more dependent of cell differentiation state and cell division rate (Supplementary figure 8a).

### Core histone turnover distinguishes dividing cells from non-dividing cells

As the turnover of core histones seemed to be related to cell division speed, we wondered whether we could distinguish cells undergoing mitosis using this metric. For each cell population corresponding to a particular time point, we plotted the distribution of L/H SILAC ratios of core histone H4, which was detected in most of the cells (Figure 7a). While iPSCs showed a single distribution suggesting a single population with fast turnover, differentiated cells gradually exhibited a second distribution that appeared following differentiation and taking over as cells further differentiated becoming the most abundant starting from TP4. The two cell populations showed very similar relative turnover between time points suggesting that the degree of differentiation had less impact than the cause of the presence of these two populations. As cells divide covering more of the culture dish and reach terminal differentiation many of them stop dividing and the presence of the two populations in plated EBs but not in iPSCs, which are actively dividing strongly suggest that these population represent dividing and non-dividing cells. The correlation between H4 and other histones of the core complex was also very high within each time point (Figure 7b) reinforcing this conclusion. Cells from the fast histone turnover population have higher levels of cell cycle-regulated proteins and higher levels of histones on average compared to the slow histone turnover population showing that they accumulate these proteins more. Importantly, there were no such distributions when plotting the abundance of histones showing that it is a property unique to histone turnover (Supplementary Figure 8b). The two cell populations can be separated with a very clear L/H SILAC ratio cutoff and thus SC-pSILAC can distinguish cells that are actively dividing from non-dividing cells. This surprising finding is very relevant in cancer research where dormant cancer cell populations induce relapse and metastasis and are less affected by chemotherapy treatment targeting fast dividing cells or avoiding immunotherapies. Our data suggests that SC-pSILAC could aid in the study of these cell populations by not only detecting and pinpointing these cells in a population but also providing abundance and turnover measurements of their proteome^41–43^. This can also have implications in stem cell research where studying re-activation of cell division of tissue-resident stem cells such as hematopoietic stem cells while maintaining cell stemness is key to advancing therapies^44–46^ and requires new analytical means.

**Figure 7.**
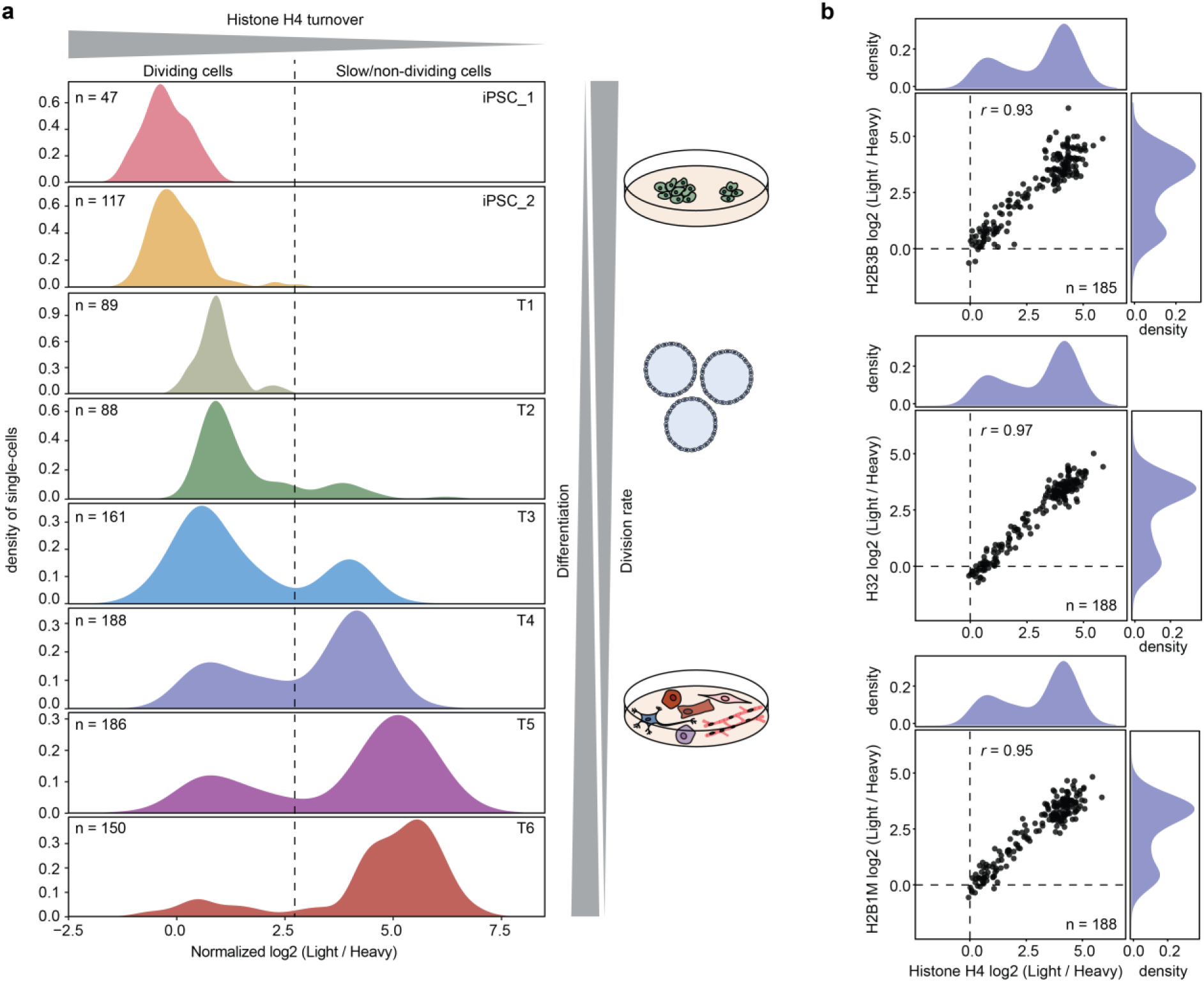
Histone turnover can differentiate dividing from non-dividing cells. (**a**) Distribution of histone H4 normalized relative turnover (light / heavy SILAC) in each cell populations chronologically from top to bottom. (**b**) Scatter plot of histone H4 normalized relative turnover (x-axis of each scatter plot) against the normalized relative turnover of histones H2B3B, H32 and H2B1M (from top to bottom). The distributions on top and right side of each scatter plot represent x and y-axis, respectively. The correlations were calculated using Pearson analysis.

## Discussion

We have performed a proteome-wide analysis of protein turnover in single cells employing a pSILAC approach, capitalizing on the recent technological improvements in SCP^33^. Our workflow includes the use of the recently introduced Evo96 proteoCHIP, which increased the analytical depth in SC-pSILAC analyses, comparable to what we have previously shown for label-free SCP analysis^33^. High sensitivity is particularly relevant for SC-pSILAC analysis as the MS signal is split between the two SILAC labels reducing the number of identified proteins.

While our experimental design precludes the measurement of protein synthesis or degradation rates because due to the lack of a reference point using a third label in non-pulsed cells and of multiple sampling intervals, this is anyway irrelevant. Indeed, every single cell is considered an independent sample, which makes it conceptually impossible to determine protein synthesis or degradation rates with current approaches, including ours.

Although protein levels do not always correlate well with mRNA levels^47^, particularly when cells are stressed (e.g. drug response, differentiation…), it is often assumed that transcriptomics and proteomics provide analogous biological insight. Since transcriptomic analyses so far have provides higher throughput and coverage than proteomics particularly at the SC level, it is the gold standard for single-cell analysis at the moment. Thus, SC-pSILAC represents a unique avenue for SCP and, in addition to recent studies highlighting PTM measurements^23,33^, it is unique by providing a completely orthogonal measurement that cannot be approached by current competing technologies.

Beyond its primary application in measuring protein turnover, SC-pSILAC offers potential in mitigating batch effects arising from sample preparation as the incorporation of the labeled amino acids is metabolic and solely dependent on the state of the cells. Additionally, the ratio of the two SILAC labels is less sensitive to cell size for proteins that are confidently detected, and therefore, does not require extensive normalization. This is particularly relevant in studies involving highly heterogenous cell populations such as cancer cells or stem cell differentiation. A drawback of SC-pSILAC similar to bulk SILAC analysis, is that it cannot be used for clinical samples. SC-pSILAC, can however be applied to organism that allow incorporation of the amino acids such as mouse^48^ and fly^49,50^. Another challenge lies in the difficulty of incorporating a third SILAC label, as the spacing between SILAC peptide precursor triplets decreases. This increases the likelihood of detecting multiple SILAC-labeled versions of the same peptide in the same DIA window, potentially compromising MS2 identification and quantification accuracy^29^. Although pSILAC is a multiplexed “labeled” measurement approach, it does not increase the overall analysis throughput. Multiplexing pSILAC experiments would necessitate a separate labeling strategy involving distinct isotopes for different cell populations, similar to what was done on bulk samples^51^. Furthermore, the incorporation of the pulsed label should be as close to 50% as possible, to facilitate the quantification, as discrepancies in signal intensity may pose challenges in quantifying labels with low intensity, thereby increasing missing values. Nevertheless, the analysis of abundance and turnover changes in CTNB1 upon CHIR99021 treatment represents one of the few functional SCP studies to date, paving the way for further functional and multidimensional measurements of protein properties in SCs.

Finally, our study of stem cell differentiation exemplifies the usefulness of such an approach in a much more biologically complex system, where obtaining multidimensional information on protein chemistry is key to unravelling new biological mechanisms. For instance, being able to directly pinpoint dividing and non-dividing cells based on the core histone turnover dynamics with no additional sorting or specific labeling holds great potential for future studies in cancer research and regenerative medicine. In addition, our study demonstrates that SCP is already a mature enough technology to envision larger scale analysis yielding relevant biological information, especially with the latest developments in analysis throughput. This dataset can be further explored to study the relationship between protein abundance and turnover to cell size, cell state and cell division rate.

## Material and methods

### Cell culture

HeLa, HEK 293T and hFF cells (ATCC) were grown in DMEM supplemented with 10% fetal bovine serum (FBS), 2 mM L-glutamine and 100 units per milliliter penicillin/streptomycin.

Human iPSCs hi12^40^ were maintained on laminin521-coated dishes in E8 medium supplemented with 0.1% BSA.

All cells were maintained in a humidified incubator (Thermo Fischer Scientific) at 37℃ and 5% CO_2_.

### EB induction

EB induction was performed as previously described^40^. Hi12 cells were dissociated from using TrypLE Express, for 5 minutes followed by scraping and resuspended in NutriStem (Saveen Werner) with 10 µM of Y-27632 (a ROCK inhibitor) and cultured in suspension in low adhesion flasks. After a week, EBs were plated on gelatin-coated 6-well plates in DMEM medium (GIBCO) supplemented with 20% (vol/vol) fetal bovine serum (GIBCO), 2 mM l-glutamine and 1% (wt/vol) nonessential amino acids (GIBCO). Medium was replaced with fresh medium every three days.

### SC-pSILAC analysis of drug treatments

For regular pulsed SILAC analysis, HeLA, HEK 293T, hFF were grown in medium containing the non-isotopically labeled version of lysine and arginine. 18 hours prior to the sorting, the medium was replaced with pre-warmed DMEM for SILAC medium (Fischer Scientific) supplemented with heavy L-lysine and L-arginine (Cambridge Isotope Laboratories) and 10% dialyzed FBS (ThermoFischer Scientific).

For the experiment with bortezomib and cycloheximide, cells were treated with two concentrations of the drugs, 1 and 10 µM and 2 and 20 µM, respectively. The drugs were diluted into the DMEM for SILAC medium at the corresponding concentrations and added to the cells to start the treatment at the same time as the pulse.

For the SC-pSILAC analysis of stem cell differentiation, medium was replaced 24h prior to sampling time by either DMEM-F12 for SILAC medium (Fischer Scientific) supplemented with heavy L-lysine and heavy L-arginine as well as E8 supplements (Gibco) and 0.1% BSA for iPSCs and non-plated EBs and with DMEM for SILAC medium (Fischer Scientific) supplemented with heavy L-lysine and heavy L-arginine and 20% dialyzed FBS for plated EBs. Cells were sorted with one week interval between each time point apart for TP6 which was sampled 10 days after TP5.

### Immunofluorescence

For immunofluorescence studies, hi12 cells and three weeks old embryoid bodies (EBs) were cultured in glass-bottom cell culture treated 96-well plate wells precoated with laminin-521 and gelatin, respectively. Cells and EBs were fixed with 4% (wt/vol) paraformaldehyde, permeabilized with 0.1% (vol/vol) Triton-X, and blocked with 10% (vol/vol) fetal bovine serum (FBS; GIBCO Invitrogen) in PBS containing 0.1% (vol/vol) Tween-20 (Sigma-Aldrich) for 1 h. Cells were incubated with primary antibody for 1.5 h at room temperature, and with secondary antibody and DAPI (Molecular Probes) for 40 min. Between incubations, specimens were washed with 0.1% (vol/vol) Tween-20 in PBS buffer three to five times. Specimens were preserved in fluorescence mounting medium (Dako), and pictures were made using a confocal fluorescence microscope (Stellaris 5, Leica). The pictures were analyzed using ImageJ2 (version 2.14.0).

### Antibodies

Anti-OTX2 (HPA000633, Atlas Antibodies, Sweden); Anti-SALL4 (HPA015791, Atlas Antibodies, Sweden); Anti-TUBB3 (AMAb91395, Atlas Antibodies, Sweden); Anti-NES (AMAb90556, Atlas Antibodies, Sweden); Anti-Oct-3/4 (AF1759, Biotechne); Anti-AFP (MAB1368, Biotechne); Anti-Actin, α-Smooth Muscle antibody (A5228, SIGMA).

### Cell sorting

Cells were detached using trypLE select (Gibco), fresh medium was added to dilute the tryplE and cells were centrifuged at 400 x g for 3 min then washed twice with Phosphate Buffered Saline (PBS) (Gibco) with centrifugation in-between, before being resuspended in degassed PBS with thorough pipetting. The cell concentration was adjusted to ∼200 cells/μl with degassed PBS. 1 cell, 10 cells or 20 cells were sorted into either label-free, LF-48 or Evo96 proteoCHIPs (Cellenion) onto pre-dispensed lysis and digestion buffer consisting of 0.2% n-Dodecyl-β-D-Maltoside (DDM), 100 mM TEAB, 10 ng/µl lysyl endopeptidase and 20 ng/µl trypsin^52^. For each condition a bulk sample was also collected by taking 1 µl of cell suspension and mixing it with 1 µl of lysis and digestion buffer. The digestion was conducted at 50℃ for 1.5 hours and quenched by adding 4 µl of 0.1% formic acid to the wells. The oil present on the plate was frozen by placing the chips in the fridge and then transferred on ice. Lastly samples were loaded onto pre-equilibrated Evotips (Evosep) following manufacturer’s protocol and stored in the fridge prior to LC-MS/MS analysis.

### LC-MS/MS analysis

Evotips containing the samples were loaded on an Evosep One (EvoSep Biosystems) LC system connected to an Orbitrap Astral MS (ThermoFischer Scientific). Samples were analyzed in 40SPD (31-min gradient) using a commercial analytical column (Aurora Elite TS, IonOpticks) interfaced online using an EASY-Spray™ source. The Orbitrap Astral MS was operated at a full MS resolution of 240,000 with a full scan range of 380 − 980 m/z when stated. The full MS AGC was set to 500% or 30 ms. MS/MS scans were recorded with 4 Th isolation window, 6 ms maximum ion injection time.

MS/MS scanning range was from 380-980 m/z were used. The isolated ions were fragmented using HCD with 27% NCE.

### Protein quantification

Protein identification and quantification was performed on Spectronaut version 18 with mostly default settings, except that “channel1” was added with no modification and “channel2” with Lys8 and Arg10 and that carbamidomethylation was not included in the fixed modifications. Exclusion of interfering fragments is added by default and cross-run normalization was removed.

## Supplementary figures

**Supplementary Figure 1.**
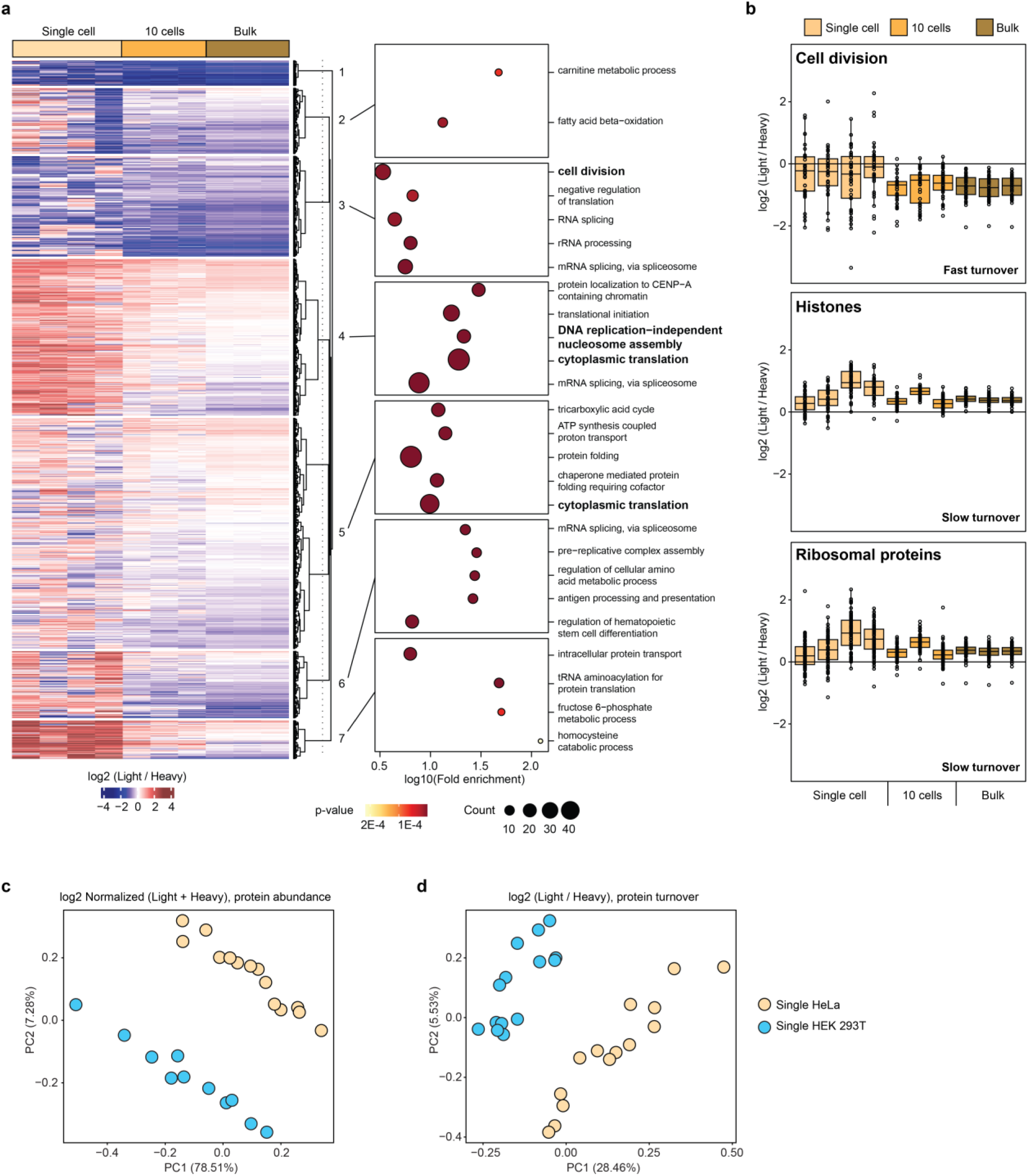
Pathways analysis of pSILAC comparison for bulk, 10 and single HeLa cells. (**a**) Unsupervised hierarchical clustering using canberra and ward.D2 methods of proteins log 2 SILAC ratio of light label divided by heavy label depicting protein turnover (left). GO enrichment analysis using DAVID of the protein present in each cluster (right). The top 5 enriched pathways according to Benjamini-Hochberg-corrected p-value are highlighted. (**b**) Turnover value for each protein involved in cell division, ribosomal proteins or histones. Horizonal line in the boxplots represent the median, 25th and 75th percentiles and whiskers represent measurements to the 5th and 95th percentiles. (**c**) PCA of single HeLa and HEK 293T cells using the log2 normalized sum of the two SILAC labels representing protein abundance. (**d**) PCA of the same samples but with the log2 relative turnover of light divided by heavy. n=4 single HeLa top, n=3 10 HeLa, n=3 bulk HeLa, n=14 single HeLa bottom, n=11 single HEK 293T.

**Supplementary Figure 2.**
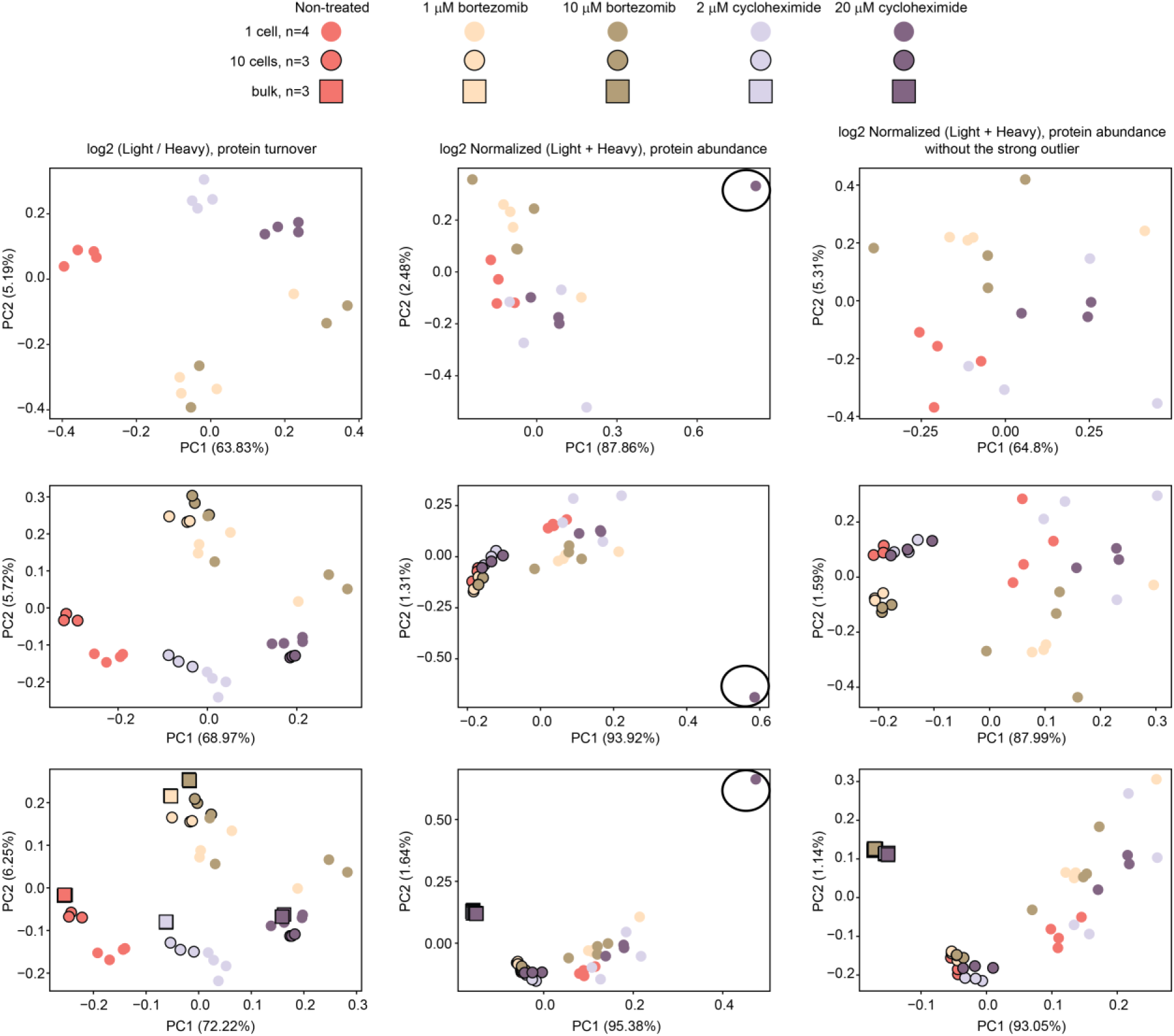
PCA of protein relative turnover and abundance in bulk, 10 and single HeLa cells non-treated and treated with bortezomib and cycloheximide. First column represents protein turnover as the log2 ratio of ligh/heavy SILAC label, second column represents protein abundances as the log2 normalized sum of the two labels and third column is the same with the strong outlier for cycloheximide treatment excluded. First row represents SCs only, second row represents single cells and 10 cell samples, third row represents SCs, 10 cell and bulk samples. N=3 for bulk and 10 cells samples, n=4 for SC of each treatment.

**Supplementary Figure 3.**
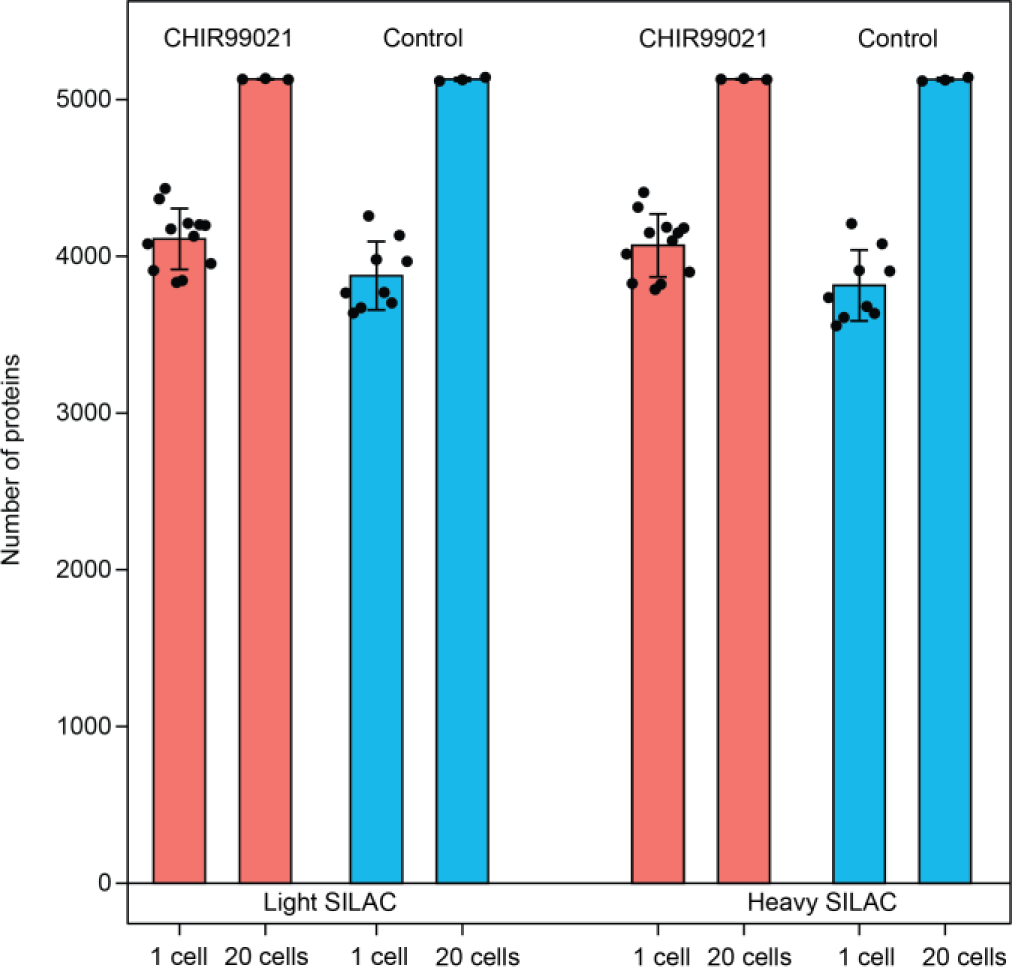
Number of proteins quantified in HeLa treated with CHIR99021 and control. Error bars represent ± the standard deviation of the mean. n=12 for CHIR99021 treatment and n=9 for DMSO-treated control.

**Supplementary Figure 4.**
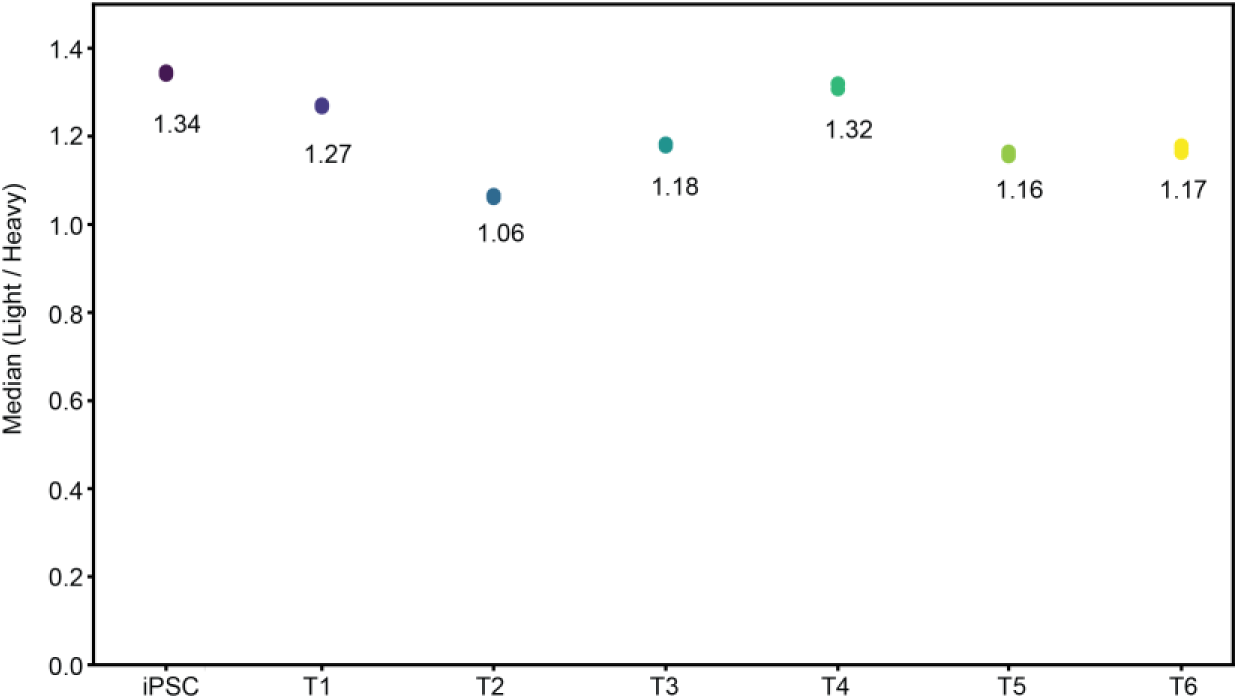
Heavy SILAC incorporation in human foreskin fibroblasts (hFF) at each corresponding sampling point of the stem cell differentiation analysis. The mean between triplicates at each time point is displayed.

**Supplementary Figure 5.**
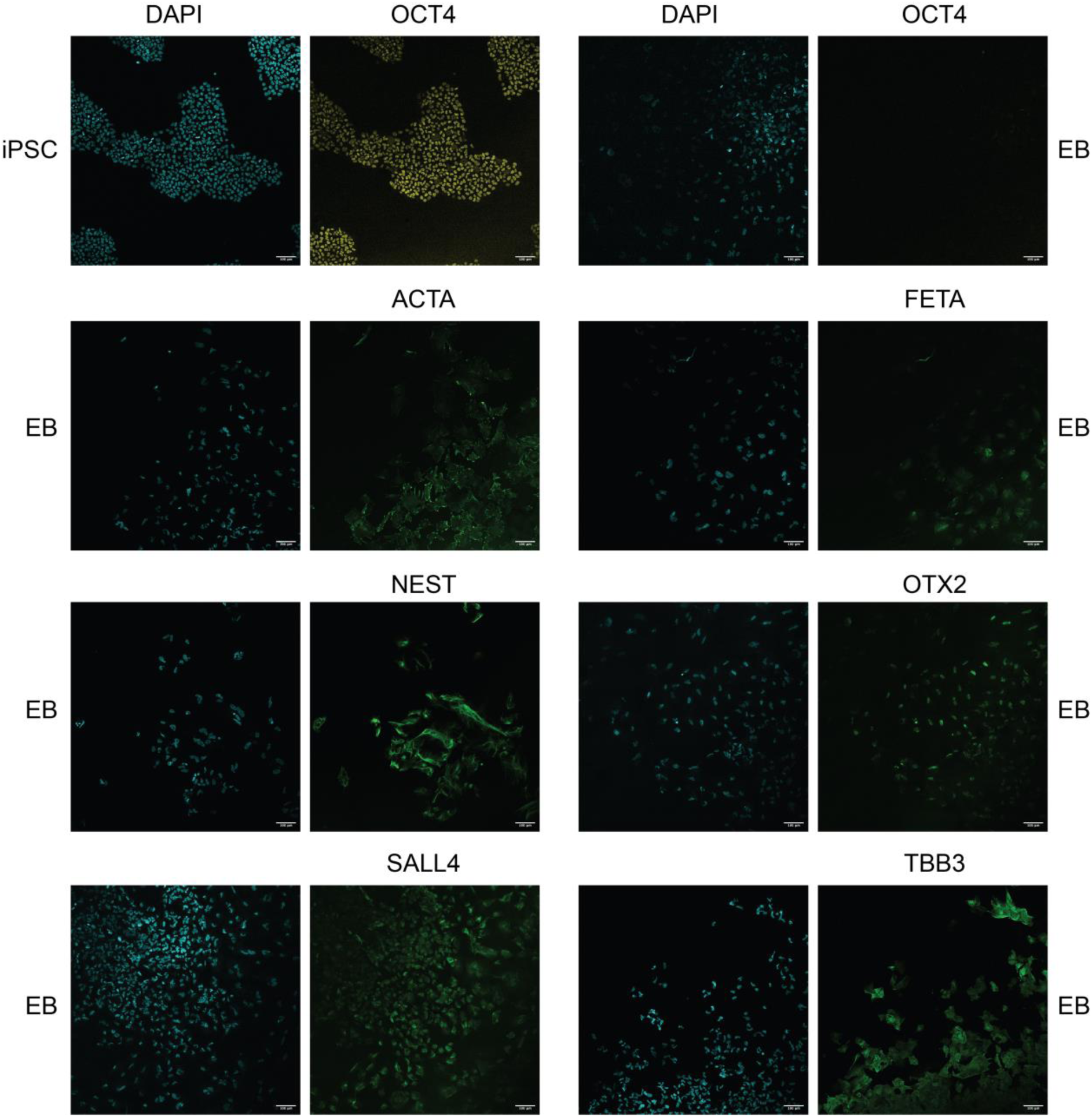
Fluorescence microscopy analysis of pluripotency and lineage markers. Each iPSC and EB samples were stained with DAPI and with antibodies directed against OCT4 for iPSCs, and OCT4, ACTA, FETA, NEST OTX2, SALL2 and TBB3 for EBs. Scale bar represents 100 µm.

**Supplementary Figure 6.**
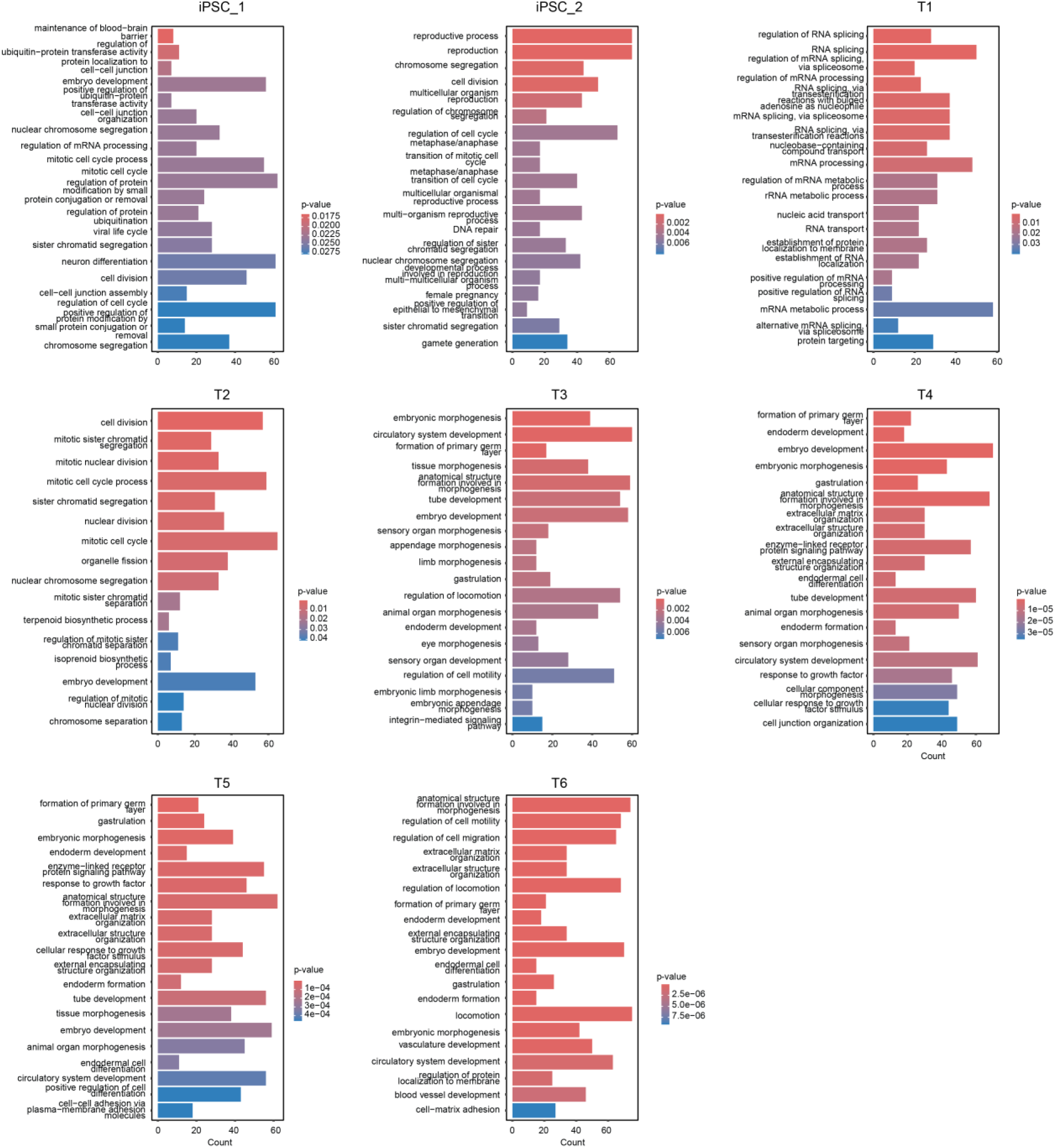
Pathways displaying fast turnover during differentiation. GO enrichment analysis employing clusterProfiler R package^53,54^ using the median normalized L/H ratio of each of the 500 protein with the lowest L/H ratio (fastest turnover) at each time point. P-values were adjusted by Benjamini-Hochberg procedure.

**Supplementary Figure 7.**
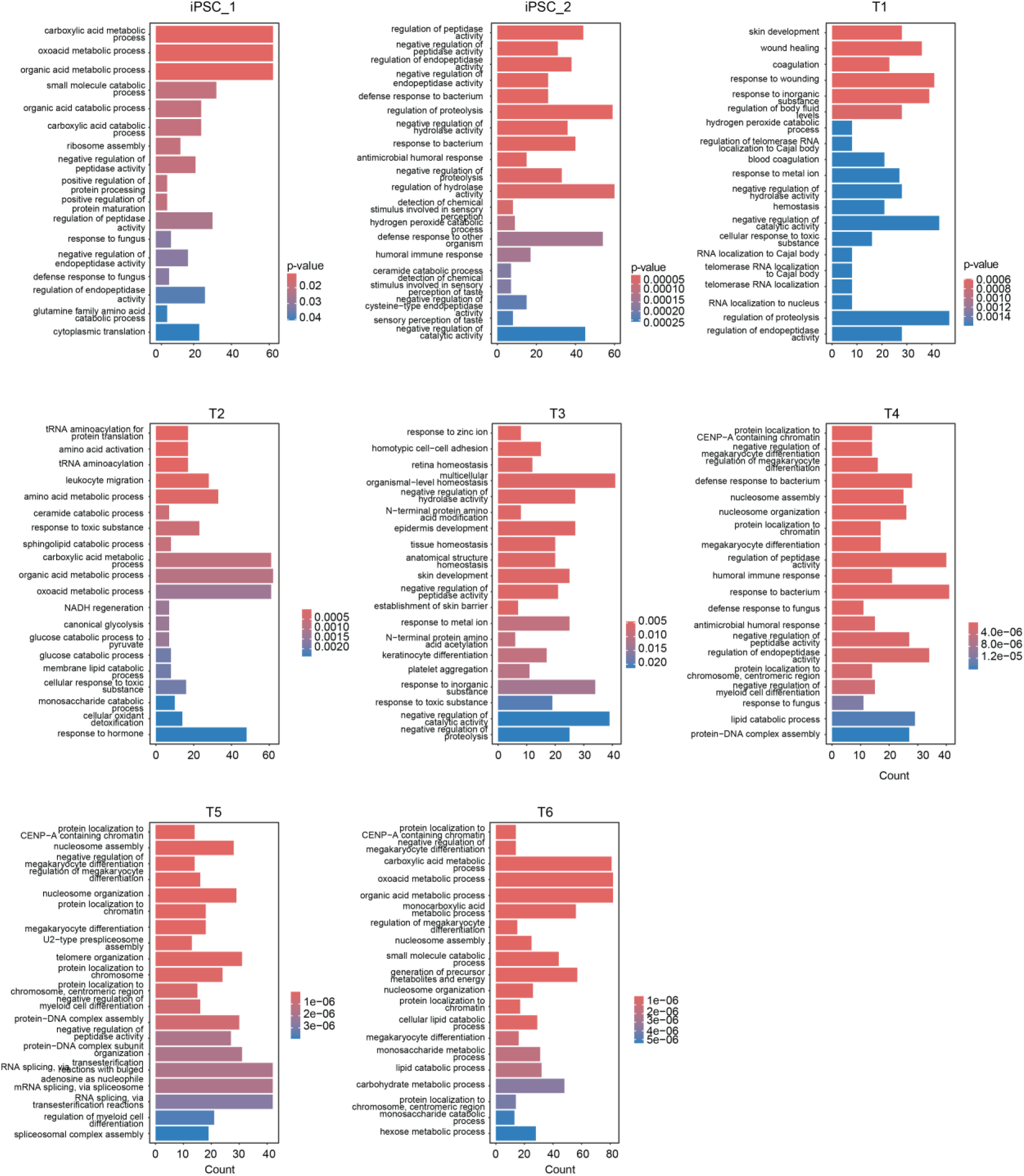
Pathways displaying slow turnover during differentiation. GO enrichment analysis employing clusterProfiler R package^53,54^ using the median normalized L/H ratio of each of the 500 protein with the highest normalized L/H ratio (slowest turnover) at each time point. P-values were adjusted by Benjamini-Hochberg procedure.

**Supplementary Figure 8.**
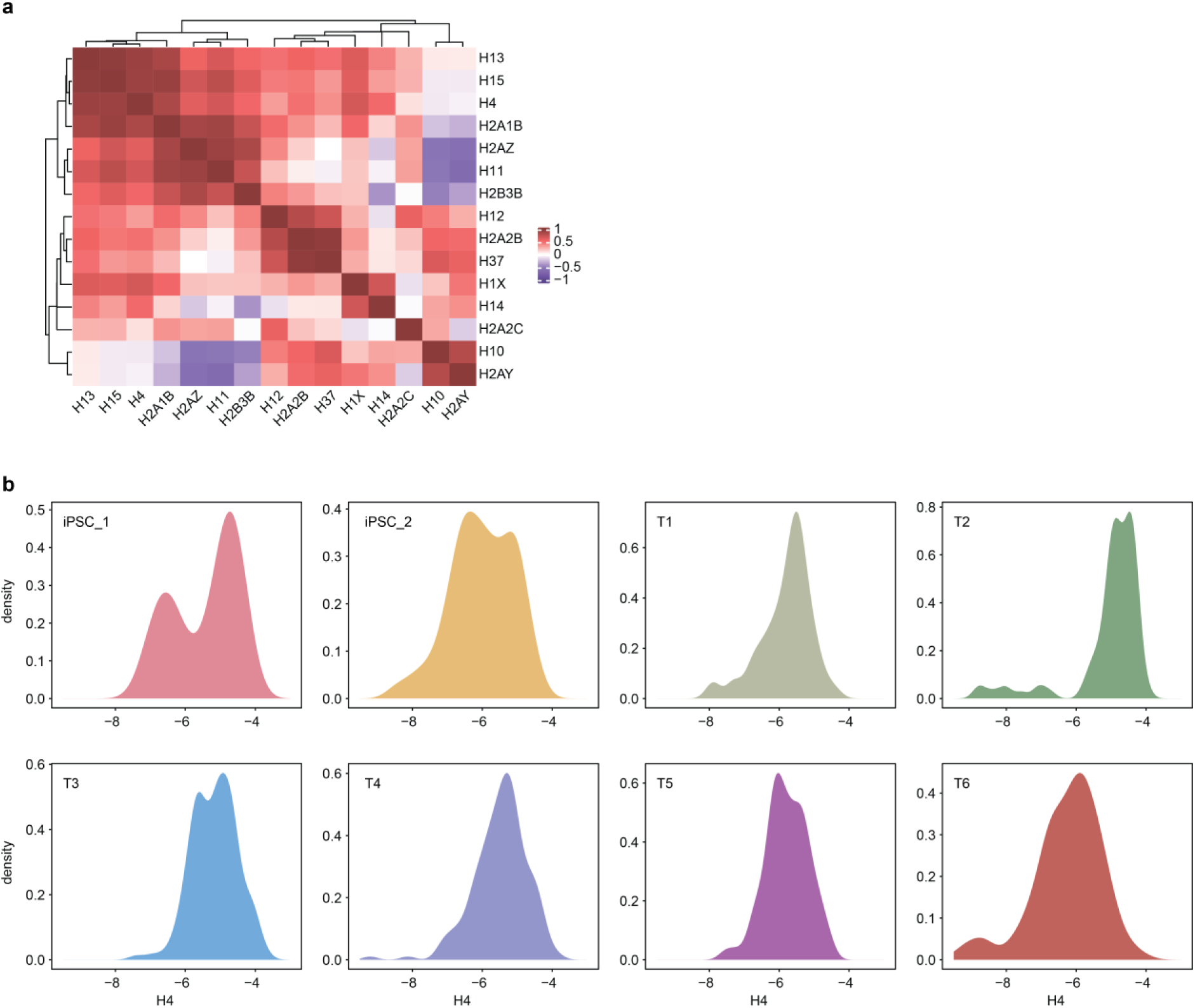
Correlation of histone protein abundance and distribution of H4 abundance. (**a**) Unsupervised hierarchical clustering using canberra and ward.D2 methods of the Pearson correlation of the abundance of each histone subunit detected in more than half the cells present in the dataset, the median value at each time point was used to calculate the correlation. (**b**) Distribution of the normalized abundance of histone H4 at each time point.

